# Accelerating leaf area measurement using a volumetric approach

**DOI:** 10.1101/2021.07.03.451015

**Authors:** Abbas Haghshenas, Yahya Emam

## Abstract

Despite the advances in the techniques of indirect estimation of leaf area, the destructive measurement approaches have still remained as the reference and the most accurate methods. However, even utilizing the modern sensors and applications usually requires the laborious and time-consuming practice of unfolding and analyzing the single leaves, separately. In the present study, a volumetric approach was tested to determine the pile leaf area based on the ratio of leaf volume divided by thickness. For this purpose, the suspension technique was used for volumetry, which is based on the simple practice and calculations of the Archimedes’ principle. The results indicated that the wheat volumetric leaf area (VLA), had an approximately 1:1 correlation with the conventionally measured optical leaf area (OLA). Exclusion of the midrib volume from calculations, did not affect the estimation error (i.e. NRMSE less than 2.61%); however, improved the slope of the linear model by about 6%. The error of sampling for determining the mean leaf thickness of the pile, was also less than 2% throughout the season. Besides, a more practical and facilitated version of the pile volumetry was tested using a Specific Gravity Bench (SGB), which is currently available as a laboratory equipment. As an important observation, which was also expectable according to the 3D shape of the leaf (i.e. expansion in a 2D plane), it was evidenced that the variations in the OLA exactly follows the pattern of the changes in the leaf volume. Accordingly, it was suggested that the relative leaf areas of various experimental treatments may be compared directly based on the volume, and independently of leaf thickness. Furthermore, no considerable difference was observed among the OLAs measured using various image resolutions (NRMSE < 0.212%); which indicates that even the current superfast scanners with low resolutions as 200 dpi may be used for a precision optical measurement of leaf area. It is expected that utilizing the reliable and simple concept of volumetric leaf area, based on which the measurement time may be independent of the sample size, facilitate the laborious practice of leaf area measurement; and consequently, improve the precision of the field experiments.

## 1. Introduction

The two-dimensionally extended organ of photosynthesis has long been the focus of plant scientists. Accordingly, leaf area is probably the most frequently measured phenotypic feature of crop canopies, which has also led to development of a relatively diverse methodology. Although at the present, various non-destructive techniques and tools have been introduced for estimation of leaf area and its twin concept of leaf area index (LAI; e.g. see Watson, 1947; Neumann et al., 1989; Weiss et al., 2004; Behera et al., 2010; Liu et al., 2010; Viña et al., 2011; Zhao et al., 2011; Confalonieri et al., 2013; Mu et al., 2017., Yan et al., 2019; and Zhao et al., 2019), the destructive approaches have remained as the most accurate (Jonckheere et al., 2003) and the direct method of leaf area measurement. Indeed, even the parameters of the non-destructive techniques are calibrated or validated based on the destructive methods, which may be a laborious practice.

Gregory (1921) is known as the first researcher who reported the measurement of leaf area in 1917. He used a ruler and celluloid protractor for *in situ* estimation of leaf area based on the leaf dimensions and 2D shape. Over the next century, various innovative approaches have been used to facilitate leaf area measurement; for instance, based on leaf weight (i.e. gravimetric method; e.g. see Watson, 1937; Huang et al., 2019a), leaf water content (Hughes et al., 1970), length and width (Cho et al., 2007; Shabani & Sepaskhah, 2017; Shi et al., 2018), or utilizing planimetric (Wolf et al., 1972) and image processing techniques (e.g.Fladung, 1991), which eventually has reached to the era of smartphone apps (Easlon et al., 2014; Schrader et al, 2017; Liu et al., 2019; Müller-Linow et al., 2019; Getman-Pickering, 2020). From the first scientific attempts for directly estimation of leaf area to the modern techniques, often there has been a need for flattening the single leaves and analyzing them separately; the practice which is time consuming and laborious, despite that the high-speed sensors or equipment might have been utilized. As a consequence, the error of operator in separating the leaves or controlling the overlaps may also arise; e.g. during area measurement of the dense sample of wheat leaves harvested before stem elongation. Quantification of such practical errors has been neglected in the literature.

The only type of destructive methods which has been used for estimating the area of a leaf pile at once, are the approaches developed based on the leaf weight. Although strong relationships between leaf area and dry and/or fresh weights have been reported (e.g. see Huang et al., 2019b), generalization of the resulted models into other genotypes or conditions may be challenging and require further studies for adjusting the parameters. Indeed, leaf weight has not a direct or intrinsic mathematical relationship with leaf geometry, and may be affected by variations in the leaf water content (particularly in the case of using fresh weight), genotype, and environmental conditions (e.g. see Niklas et al., 2009; Weraduwage et al., 2015; and Lin et al., 2018). Therefore, accurate estimation of leaf area according to the weight, requires model calibration. Moreover, determination of dry weight also requires additional time and equipment for oven drying of the samples.

In contrast to the weight, leaf volume is a direct contributor to the leaf geometry; and the 2D area may be calculated simply through dividing the leaf volume by its thickness. Therefore, the resulted area may be independent of the growth condition, genotype, water content, or other variables. Consequently, the focus of the present study was on the idea of using this equation for facilitating the direct measurement of leaf area. Huxley (1971) measured the volume of a single leaf with precision of ± 0.01 ml, using an innovative volumeter which was made based on liquid displacement. Also utilizing the Archimedes’ principle, Hughes (2005) introduced a modified version of hydrostatic weighing, i.e. the suspension technique, for precision volumetry of small objects. He reported that the new method was comparatively more accurate and reproducible than the other conventional methods developed based on water displacement. Besides the high precision, the suspension technique is simple and fast, as in practice, the volume can be measured only by weighing the object out and under water.

The purpose of the present study was evaluating the option of calculating leaf area based on the ratio of leaf volume divided by thickness; which might accelerate the operation by making it possible to accurately and simultaneously measure the volume of a leaf pile. Considering the relatively high frequency and importance of wheat leaf area measurements in crop science, the focus was put on this species, and also on the tillering phase, in which the conventional leaf area measurement is more challenging, due to the smaller size of the leaves usually wrapped in a dense pile.

## 2. Materials and Methods

### 2.1. Sampling

Leaf samples were collected during the 2020-21 growing season, from a production wheat field located at the School of Agriculture, Shiraz University, Iran (29° 73′ N latitude and 52° 59′ E longitude at an altitude of 1810 masl). The wheat cultivar Sirvan was planted on October 11, 2020 with plant density of 450 plants/m^2^ using a row planter. 150 kg nitrogen/ha (as urea) was applied in three equal splits i.e. at planting, early tillering, and anthesis. Field management was carried out throughout the season according to the local practices. In order to measure leaf dimensions, a 30×30 cm quadrat was used for random sampling of leaves, from tillering to flag leaf emergence. In each sampling date, leaf length, leaf width, diameter of midrib, and leaf thickness of up to 35 leaves were measured using a ruler and a 0.01 mm micrometer (Asimeto, Germany). Thicknesses of lamina and midrib were measured separately at three points throughout the leaf length: (i) immediately adjacent the leaf base, (ii) at the middle, (ii) and near the tip (because the midrib at the tip was not distinguishable easily, the thickness at 2 cm away from the tip was recorded). The average of the three values of lamina and/or midrib thicknesses was reported as the mean leaf thickness, and mean midrib diameter, respectively.

### 2.2. Leaf volumetry and volumetric leaf area

The leaf samples used for scanning and volume determination were harvested at mid-tillering. Several leaves were cut into different pieces (from <1 cm^2^ to full leaf). Then, the volume of each piece was measured separately using the suspension technique reported by Hughes, 2005; which is based on the Archimedes’ principle. For this purpose, a PVC beaker was filled with 250 ml distilled water and put on a 0.001 g weighing balance (A&D, FX-300GD, Japan). The balance was then tared. As the density of leaf is generally lower or around the density of water, a relatively heavy cage or retainer is required to keep the sample under water. Therefore, a steel paper clips was used. It was hung from a holder arm and completely immersed in water using a single line of copper electric wire (of 0.19 mm diameter), with a steel hook at the end. There was no contact between the immersed set and the wall or bottom of the container; so that the clips was completely suspended in the water. Under the stationary situation (i.e. after the clips stopped moving), the weight was recorded as Δ*W*_*r*_, which is the change in the weight recorded by the balance due to the immersion of the retainer. Notably, here the term retainer includes the clips, hook, and the underwater part of the line. Thereafter, the clips, hook, and line were taken out of the water, and dried. This measurement was repeated two or three times, to recognize the potential errors, e.g. due to the presence of tiny bubbles. To ensure that the same length of the line was placed under water in every iteration, the level of water was marked on the beaker using a thin-tip marker (which was 1 cm above the hook). Before each iteration, the level of water was controlled by adding water, and the balance was tared again. Water temperature was also measured using a mercury liquid thermometer. Then, each of the 14 single leaf samples (either a piece or full leaf) was attached to the clips, and the measurements were repeated separately. Enough care was taken to prevent or remove bubbles from the samples and clips.

According to the Archimedes’ principle, since the immersed set (retainer with/or without leaf sample) was stationary, the downward gravitational force had been balanced by the upward buoyancy and line tension (for more information, see Hughes, 2005). Indeed, the volume of the immersed set was equal to the volume of displaced water with the same size and shape. Therefore, the recorded weights (Δ*W*) were equal to the weight of displaced water, and the volume of the immersed set could be calculated directly by dividing the weight by the density of water at the recorded temperature. Accordingly, the volume of leaf sample was calculated as below:

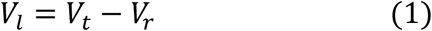

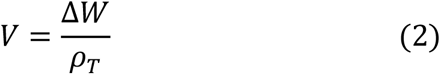

*So, based on equation 2:*

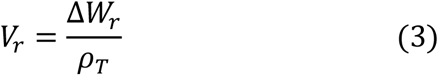

*From* *equations* 1 & 2:

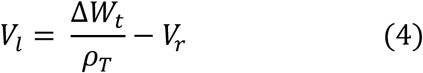

where *V*, Δ*w*, and *ρ* are volume (mm^3^), the change in the weight (exactly the values recorded by balance, g), and the water density (g/mm^3^) at *T* °C, respectively. Also, the subscripts *l, t*, and *r* stand for *leaf, total*, and *retainer*, respectively; so the *V*_*t*_ and Δ*w*_*t*_ indicate the total volume and the change in the total weight of the immersed set, which was included the leaf sample, clips, hook, and the underwater part of the line. Since a single clips was used in the experiment, the volume of retainer (*V*_*r*_) was calculated at the first step, and used in every other calculations. Values of water density at the measured temperatures were taken from *CRC Handbook of Chemistry and Physics* (Lide, 2005). In this experiment, the recorded water temperatures were in the range of 21.5 to 23 °C.

Volumetric leaf area (mm^2^) was simply calculated as:

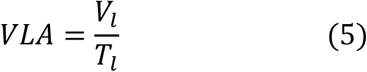

where *VLA, V*_*l*_, and *T*_*l*_ are the volumetric leaf area (mm^2^), volume of leaf sample (mm^3^), and leaf thickness (the mean lamina thickness of each single leaf, mm), respectively. In the volumetric leaf area measurement, the leaf thicknesses of the 14 samples were measured every 2 cm on the leaf length using micrometer (except for those pieces shorter than 2 cm which had only one reading; see section 2.1).

Depended on the length of the samples, volume of midrib was estimated either as the volume of a cylinder or a cone with the length equal to the length of sample. For cuts of leaves, in which the thickness of midrib shows negligible variation throughout the length, the midrib was supposed as a cylinder; while in the case of complete leaves it was taken as a complete cone with the base diameter equal to the thickness of leaf base, and a height as long as the leaf length. So:

*For small samples* (*cylindrical midrib*):

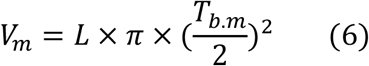

*For complete leaves* (*conical midrib*):

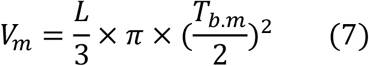

where *V*_*m*_, *L*, and *T*_*b*.*m*_ are volume of midrib (mm^3^), length of sample (mm), and basal thickness (diameter, mm) of midrib.

### 2.3. Optical leaf area measurement

Immediately after volumetry, each leaf sample was air-dried for several minutes, unfolded, and pasted on a single A4 white glossy cardboard, using a glue stick. The highest care was taken to cause complete contact between the surfaces of leaf and the cardboard, and prevent wrinkling. Then, the cardboard was scanned at various resolutions including 200, 300, 400, 600, 800, 1000, and 1200 dpi (dots per inch), using a A4 scanner (Genius ColorPage-HR7X Slim). The purpose of testing different resolutions was evaluating the effect of this factor on the precision of area measurement. Considering the 3D structure of the leaf (particularly the midrib) which had made some small bulges in the cardboard, and also for reducing the effect of environmental light, a weight of 800 g was put on the scanner cover to compress the cardboard and consequently maximize the contact between the surfaces of leaves and the scanner glass. Also, after assessing several settings of the scanner for achieving the optimum contrast between the green leaf surfaces and the white background, the brightness and contrast were set to 55% and 65%, respectively.

While the images could be processed by various methods, software or exclusive codes, they were analyzed using Adobe Photoshop CC 2017, which readily provides professional selection tools for segmentation and extracting the ROI (range of interest, i.e. green leaf area) in a reliable and simple way. Therefore, leaf area of each sample was selected with highest care, using Magic wand and other selection tools (tolerance was set to 20). Then, number of selected pixels (SP) of each sample was recorded from Histogram tool (before each reading, the “Uncached refresh button” was pressed). Finally, the area (mm^2^) of each leaf sample was calculated as below:

*Considering the square shape of pixel*:

*If* “*r*” *is the image resolution* (*dpi*), *there are* “*r*” *pixels* (*dots*) *per each inch or* 25.4 *mm of image length*/ *or width. So, we have* 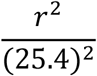 *pixels per* 1 *mm*^2^.

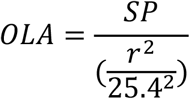

*or*:

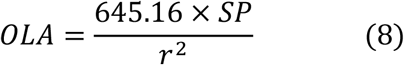

where *OLA* and *SP* are the optical leaf area (mm^2^) and number of selected pixels of each leaf sample, respectively. Here, the concepts of “*optical leaf area”* and “*volumetric leaf area*” are used for the measured leaf area by scanning, and the leaf area calculated based on the volumetric approach, respectively.

### 2.4. Utilizing Specific Gravity Bench

In order to evaluate the option of using Specific Gravity Bench (SGB) for a faster and more facilitated leaf volumetry, a SGB was equipped with a calibrated load cell (Zemic, L6D-C3-2.5kg-0.40B, China), a load cell monitor (Tika, TD-1000, Iran), and a cylindrical cage made of stainless steel (see Fig. 1). Here, the mean required time for calculating the volumetric area of 10 leaf piles were estimated. Similar to the method described in section 2.2, the volume of leaf pile was measured by the main idea of suspending the samples in the water. However, there were some minor differences in the operation and calculations. Despite the first method in which the whole system including the water container was put on the balance, in SGB technique, only the steel cage containing leaf sample was hung from the load cell. So, calculations were independent of the weight of water and the container. Similar to the first experiment, leaves were harvested at mid-tillering. Before volumetry, 5 leaves were selected randomly from each pile, their middle thicknesses were measured (one reading per leaf), and finally the mean thickness of the sample was calculated.

**Figure 1.**
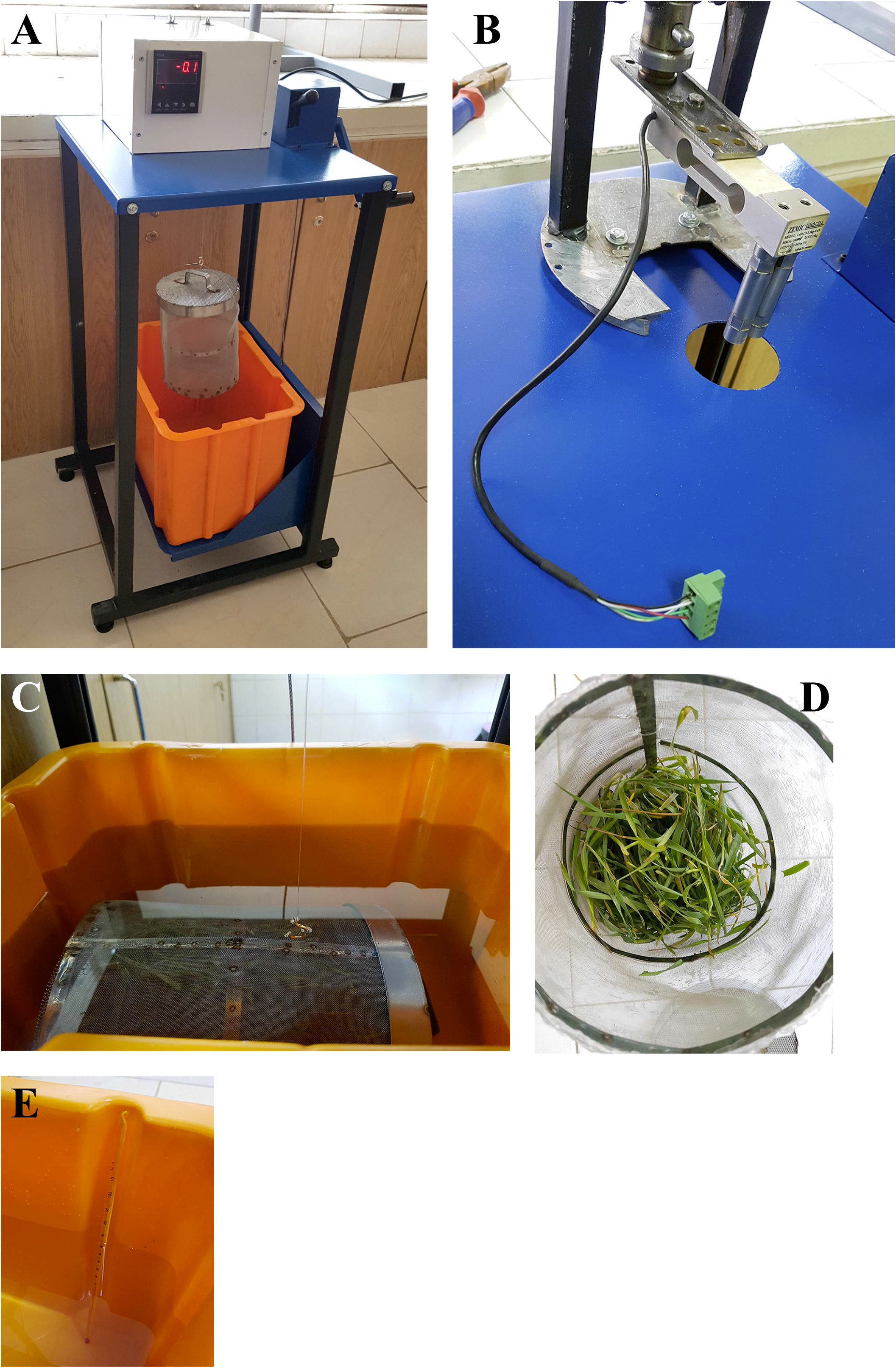
Leaf volumetry using Specific Gravity Bench (SGB). (A) An overview of the bench. The empty steel cage is hung from the load cell. (B) The SGB bench was equipped with a load cell. (C) The cage containing the leaves is immersed in the water. (D) A view of the cage containing leaves. (E) Water temperature was measured regularly using a liquid thermometer.

Similar to the previous method (Eq. 1, section 2.2), the net volume of leaf sample is equal to the total volume of the immersed set minus the retainer volume. For measuring the volume of retainer (the steel cage), the hook was hung from the load cell using a fishing line, and the load cell was tared. The weight of the dry and empty cage was recorded out of water as *W*_*r*1_. Then, it was attached to the hook, and gently immersed in water (to avoid bubbles) by turning the bench crank and raising up the water container. In the stationary situation, the weight was recorded as *W*_*r*2_. In contrast to the previous method, here a nylon ring with 2 cm diameter (made of a narrow fishing line) was fastened to the cage by which the cage was attached to the hook. Indeed, the retainer was only included the cage with a small part of the nylon ring (with a negligible volume, which also might be included in the calculations); so the hook or the holder line were kept out of the water. Based on the Eq. 3, the volume of retainer was calculated as follows:

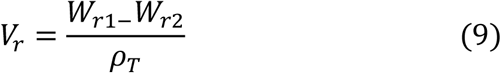

where *V*_*r*_ and *ρ*_*T*_ are the volume of retainer (mm^3^), and water density (g/mm^3^) at *T* °C. Then, the weight of each leaf pile was measured out of water and recorded as *W*_*l*1_ (if no other balance was available, the SGB load cell and its cage could be used, provided that they were dry). After determining the volume of the retainer, the leaf sample was put in the cage, and immersed in the water. The underwater weight of the immersed set was recorded as *W*_*T*2_ (“*T*” stands for *Total*). Again, it should be emphasized that for ensuring the results and removing the effect of probable bubbles, the underwater weighing (both for volumetry of the empty retainer and total set) was repeated three times by taking out the cage and re-immersing it in the water. The water temperature was also recorded regularly. Volumetric calculations for the leaf pile was as follows:

*From* *equations* 1&2:

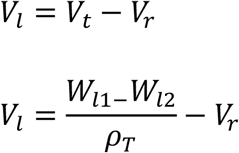

*The net underwater weight of the leaf sample is equal to*:

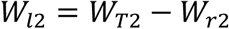

*so:*

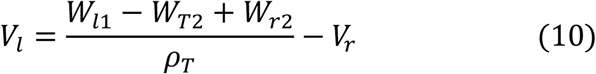

*Also using Eq. 9, a more practical form of Eq. 10 may be achieved, which is independent of W*_*r*2_:

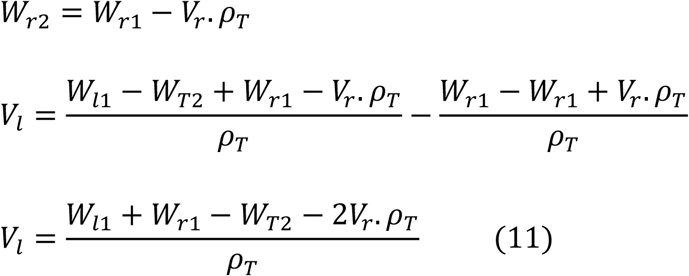

where *V*_*l*_ and *V*_*r*_ are the volumes of leaf and retainer (mm^3^), *ρ*_*T*_ is the water density (g/mm^3^) at *T* °C, and *W*_*l*1_, *W*_*T*2_, and *W*_*r*2_ are the weight (g) of leaf sample out of water, the underwater weight of the total immersed set (retainer and leaf sample), and the underwater weight of retainer, respectively. So, having the fix values of *W*_*r*1_ and *V*_*r*_, it is enough to record three parameters of *W*_*l*1_, *W*_*T*2_, and water temperature in each running of the leaf volumetry. It should be noted that *W*_*l*1_ and also *W*_*r*1_ have to be measured when the leaves and the retainer are dry; however, in the SGB approach, there is no need to dry the retainer before each measurement of the underwater weights (*W*_*T*2_).

Since the calculations could be simply carried out using formulated spreadsheets, only the duration needed for running the practice for each sample was recorded, i.e. included weighing of the sample under and out of water, recording temperature, and filling and emptying the cage. Moreover, because the volume of the retainer (*V*_*r*_) is consistent as long as a single cage is used, there was no need to repeat its volumetry.

### 2.5. Statistical analyses

In order to evaluate the effect of using mean leaf thickness of pile on the volumetric area, an additional analysis was carried out entitled Pile Analysis (PA). For this purpose, the data of the 14 leaf samples which were analyzed in sections 2.2 & 2.3, were used. These samples were virtually grouped into 100 piles of 7 samples. Then, the volumetric and the optical leaf areas of each pile were determined and compared with each other. The volumetric leaf area of each virtual pile was simply calculated by dividing the summation of the VLAs of the single samples included in the pile (section 2.2), by their mean thicknesses. The summation of OLAs (section 2.3) were also used as the optical leaf area of the pile. Grouping of the samples was carried out without replacement, using the “Data sampling tools” of XLSTAT (Version 2016.02.28451; Addinsoft). Furthermore, a similar method was utilized to assess the efficiency of sampling on the estimating mean leaf thickness of the pile. Using the data recorded in each of the last three sampling dates, 100 groups of 5 leaves were selected randomly and without replacement. Then, the mean leaf thickness estimated by sampling was compared with the actual mean of population which was the result of measuring of every 35 leaves in each date. Considering the gradient of leaf thickness from the base towards the tip of leaf, the average of 3 readings per leaf i.e. the thickness measured at base, middle, and tip of the leaf (lamina), was recorded as the leaf thickness.

All other statistical analyses were conducted using XLSTAT and IBM SPSS Statistics for Windows (Version 19.0, Armonk, NY: IBM Corp.).

## 3. Results

As shown in Fig. 2, there were very high correlations among the leaf area values measured using the 1200 dpi versus the lower resolutions. The normalized root mean square errors (NRMSE) of correlations ranged between 0.056% to 0.212%, which obviously indicates that scanning leaves even with the resolutions as low as 200 dpi could be precise enough to be used in leaf area measurements.

**Figure 2.**
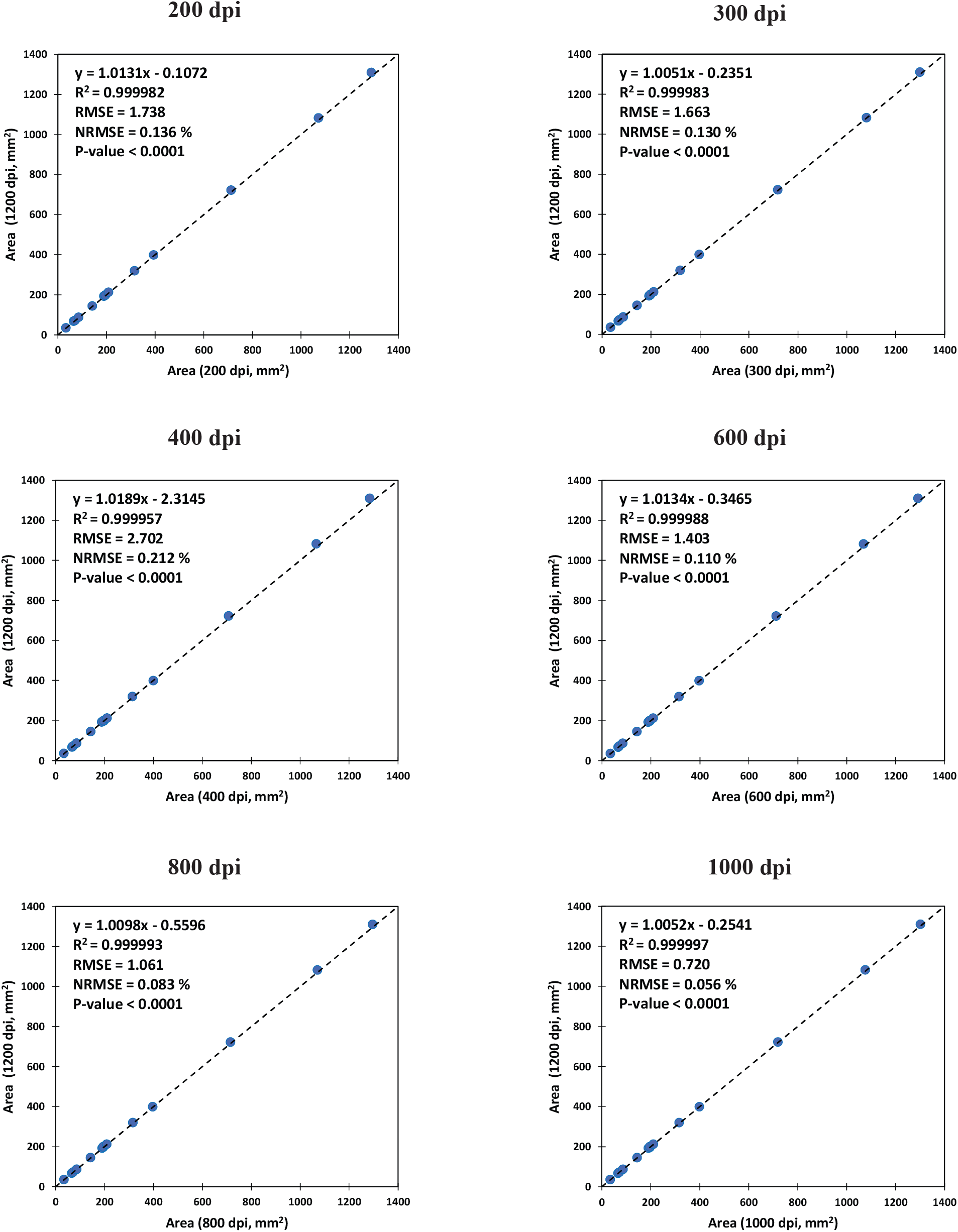
The correlation between the optical leaf area values measured by scanning with 1200 dpi (dots per inch) versus lower image resolutions.

Figure 3 indicates the correlation between optically and volumetrically measured leaf areas. In general, the results of the optical and volumetric approaches were highly correlated (NRMSE=2.61%), regardless of whether or not the volume of midrib was included in the calculations (Fig. 3A vs. 3B). However, when the volume of midrib was estimated and excluded from the calculations, the predictions were improved; i.e. the slope of the regression line became closer to 1 (0.9375 vs. 0.9903; i.e. 5.63% improvement). In average, the volume of midrib was 4.5% of the total leaf volume, with a range between 2.0% to 8.1% (data not shown).

**Figure 3.**
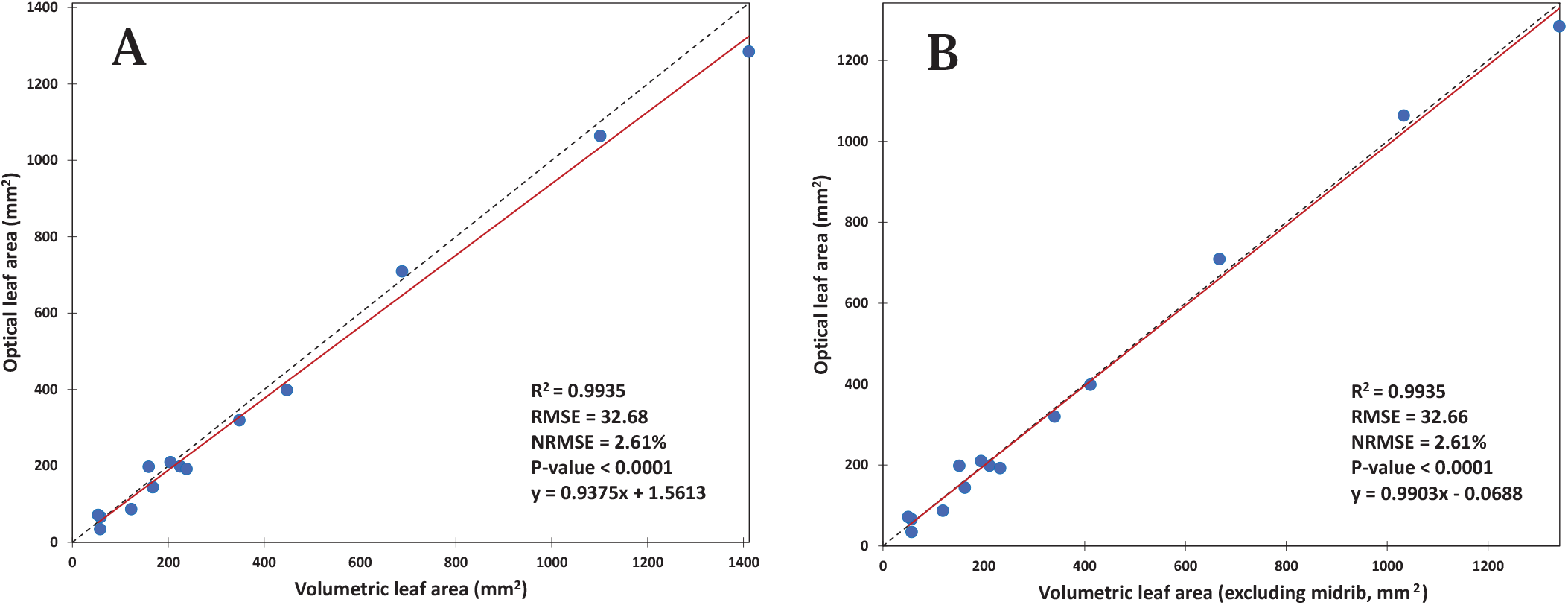
The correlation between optically and volumetrically measured leaf areas (A) including and (B) excluding the volume of midrib. NRMSE: normalized root mean square error (calculated by dividing the RMSE by the range of the optical values).

A similar trend was also observed in the Pile Analysis, where the prediction of pile leaf area was simulated using combinations of single leaves/ or leaf pieces (Fig. 4). Again, NRMSE values remained around 2%, regardless of inclusion or exclusion of midrib. By removing the share of midrib volume, the slope of the regressed line was also increased from 0.873 to 0.923 (i.e. 5.78% improvement).

**Figure 4.**
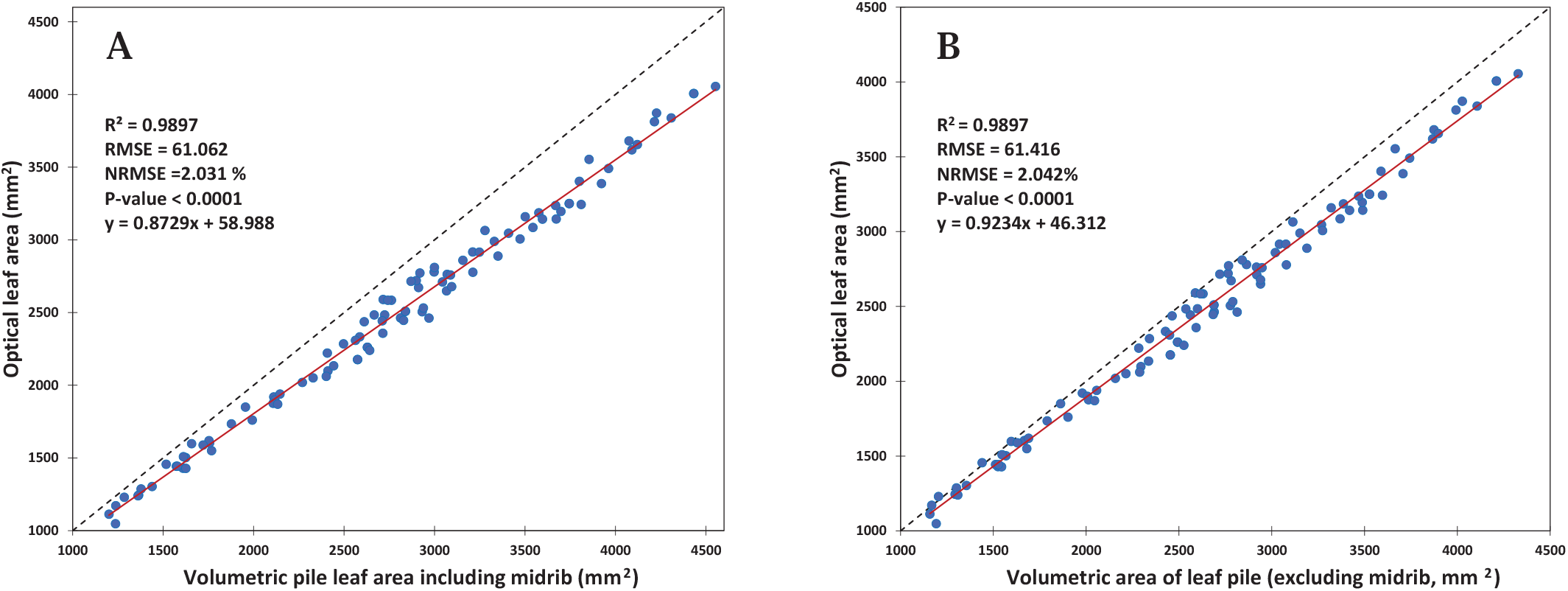
Results of Pile Analysis, i.e. the correlation between optical and volumetric areas of the leaf piles (A) including and (B) excluding the volume of midrib. Each sample was consisted of 7 leaves (or pieces of leaves) randomly chosen from the possible combinations of the 14 measured pieces (without replacement).

For evaluation of the effect of sampling and variations in the leaf thickness, several analyses were carried out. Table 1 represents the coefficients of variation (C.V.) in some properties of leaf dimension. Irrespective of the location of measuring point on the leaf, seasonal or diurnal variations in the thickness were the least, compared with the other leaf dimensions i.e. length and width.

**Table 1.** Coefficients of variation of the leaf dimensions and midrib (C.V., %).

| Days after sowing (DAS) | Leaf length | Leaf width | Base thickness | Middle thickness | Tip thickness | Mean thickness | Base midrib thickness | Middle midrib thickness | Tip midrib thickness | Mean midrib thickness |
| --- | --- | --- | --- | --- | --- | --- | --- | --- | --- | --- |
| 93 | 49.566 | 19.579 | - | 24.913 | - | 24.9135 | - | - | - | - |
| 115 | 21.763 | 18.762 | 10.732 | 18.188 | 17.482 | 12.774 | - | - | - | - |
| 136 | 7.078 | 29.737 | 16.078 | 13.331 | 13.874 | 13.465 | 20.814 | 9.795 | 18.399 | 14.139 |
| 140 | 22.493 | 14.158 | 15.014 | 11.038 | 13.803 | 11.831 | 29.730 | 12.456 | 8.469 | 17.311 |
| 143 | 18.513 | 20.055 | 11.615 | 9.586 | 13.235 | 8.828 | 23.301 | 7.150 | 9.991 | 14.290 |
| 157 | 11.219 | 11.047 | 12.836 | 8.364 | 16.527 | 10.427 | 17.638 | 7.554 | 13.357 | 11.419 |
| 161 | 13.348 | 6.052 | 11.496 | 10.656 | 14.278 | 9.438 | 16.551 | 9.623 | 11.293 | 12.147 |
| 167 | 18.228 | 17.332 | 12.607 | 12.908 | 10.729 | 9.745 | 22.430 | 13.007 | 10.224 | 12.143 |
| 174 | 19.512 | 17.564 | 14.689 | 10.305 | 13.977 | 11.784 | 23.232 | 14.576 | 11.710 | 16.781 |
| 182 | 19.607 | 14.170 | 13.050 | 15.901 | 15.649 | 13.499 | 15.203 | 15.344 | 15.519 | 13.458 |
| Mean | 20.133 | 16.845 | 13.124 | 13.519 | 14.395 | 12.671 | 21.112 | 11.188 | 12.370 | 13.961 |
| C.V. of season | 21.033 | 19.258 | 15.344 | 12.895 | 14.481 | 11.824 | 23.556 | 14.336 | 12.461 | 15.797 |

Base, middle, and tip thickness: thickness of base, middle part, and tip of the leaf. Mean thickness: The average of base, middle, and tip thickness. C.V. of season is calculated using the total data of the season.

Figure 5 represents the results of estimating the average pile thickness in the last three sampling dates, based on sampling of 5 leaves. The difference between the averaged thickness of population and the overall estimated mean of samples ranged between 1.01% to 1.62%. Since the volumetric leaf area is estimated by dividing the leaf volume by thickness, these quantities of differences may be also taken directly as the error of leaf area estimations (provided that the volumetry has been carried out perfectly).

**Figure 5.**
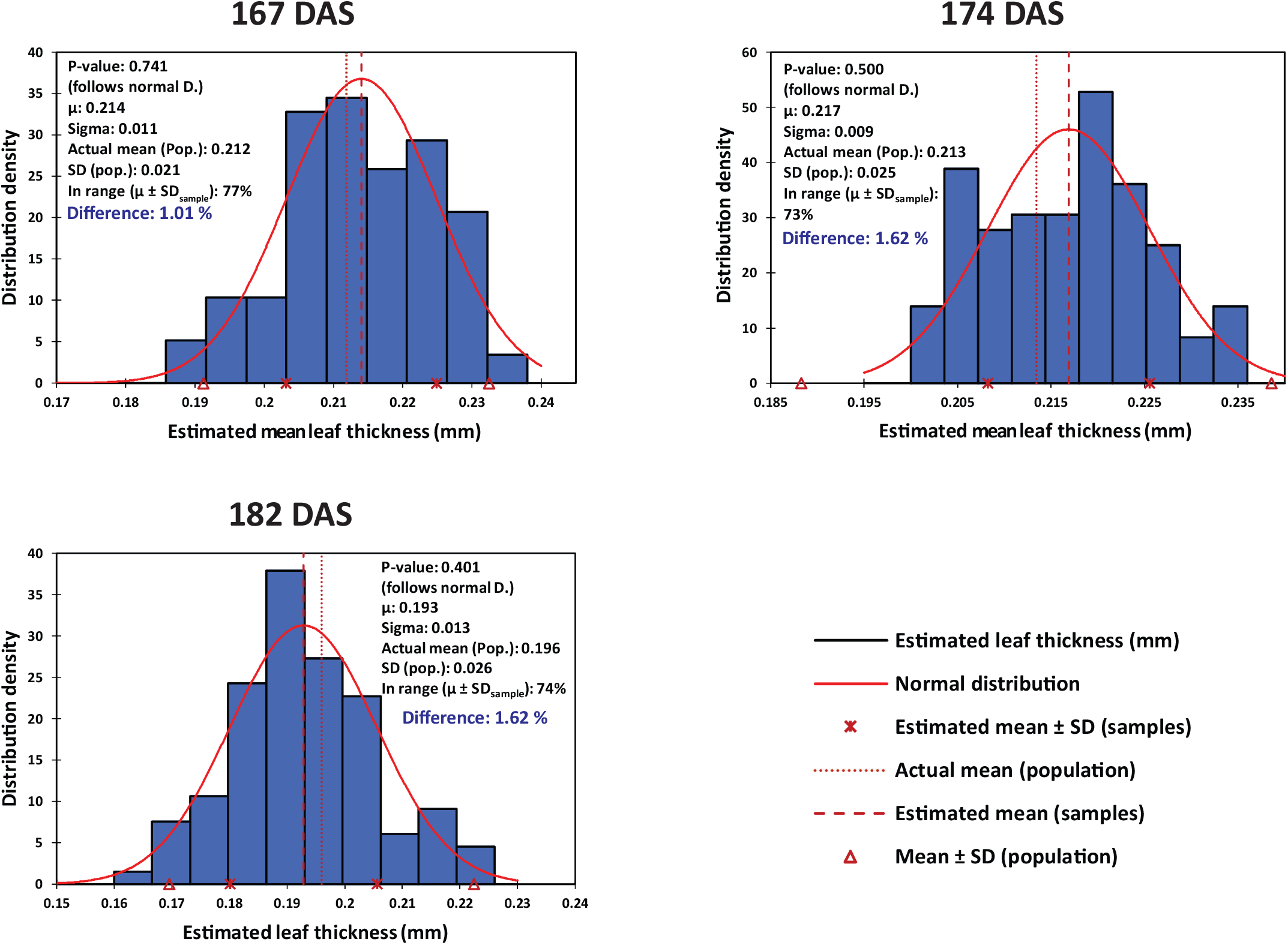
The precision of estimating mean leaf thickness of the pile, based on the sampling of 5 leaves from the piles harvested in different dates. In each date, thickness of 35 leaves were measured. Then 100 virtual groups of 5 leaves were formed randomly, and mean thickness of leaves in each group was estimated based on the average of thickness values. The bars indicate the distribution of mean thickness of the 100 groups. DAS: days after sowing; µ: overall mean thickness of piles; SD: standard deviation; Difference: the difference between the averaged thickness of population and overall mean thickness estimated by sampling.

As shown in Fig. 6, variations in the leaf area were extremely similar to the leaf volume, irrespective of the volume of midrib was included in the estimations or not. The coefficient of Pearson correlation between leaf volume and leaf area was 0.996 in both types of estimation (Fig. 6). This strong relationship may eliminate the need for measuring the leaf thickness, when the purpose is relative comparison or evaluation of variations in leaf area of treatments, rather than calculating the absolute quantitative areas.

**Figure 6.**
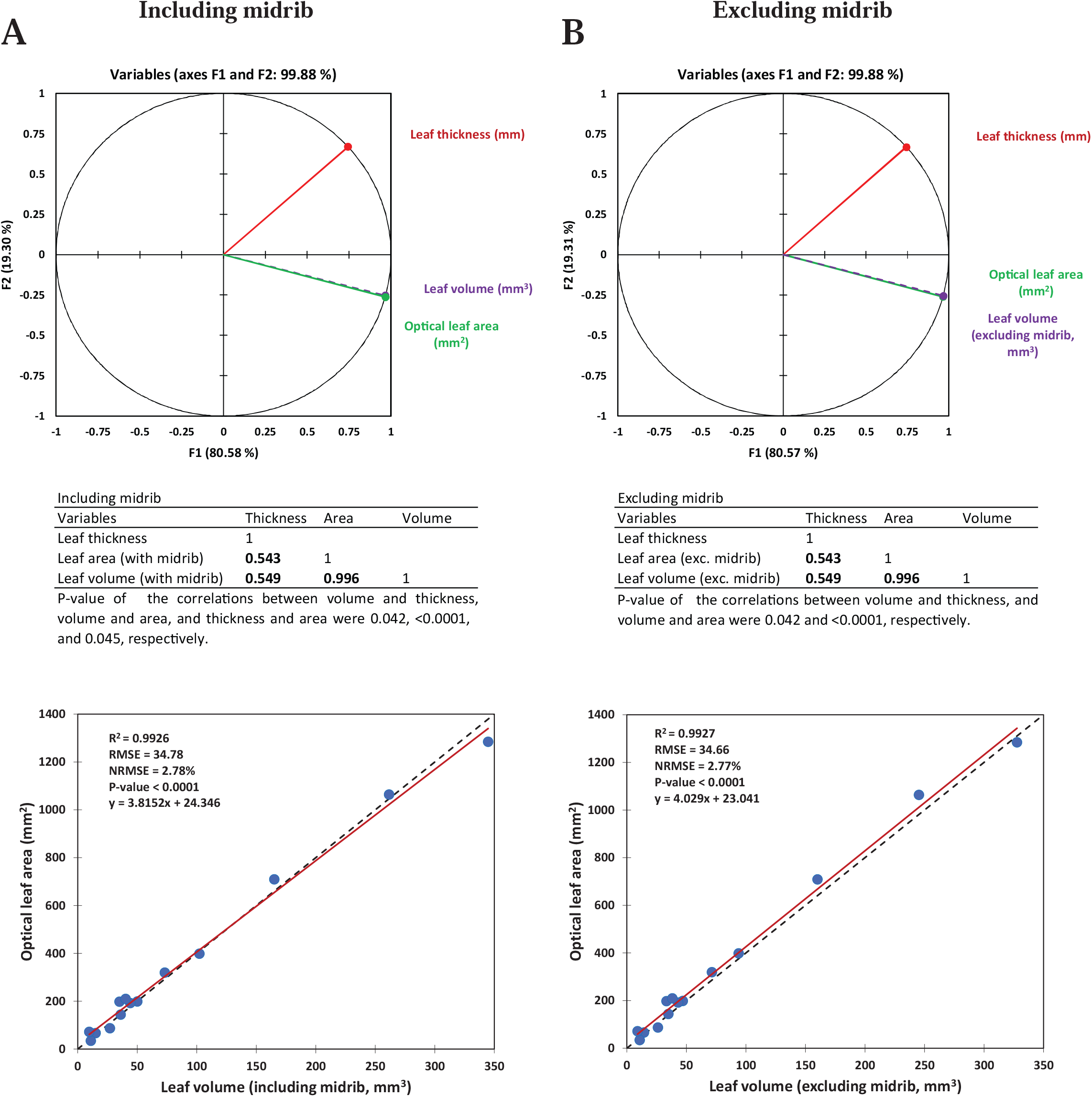
The correlation between leaf volume and optical leaf area, (A) including and (B) excluding the volume of midrib.

Moreover, Table 2 and Fig. 7 represent additional information about the dynamics of leaf dimension properties throughout the season. It was obvious that although the correlations between leaf thickness with length or width were significant, these relationships were not strong. Moreover, the mean midrib diameter (thickness) was more correlated with mean leaf thickness than leaf length or width.

**Table 2.** The correlation coefficients (Pearson) among the leaf dimensions.

| Variables | leaf length | Leaf width | Base thickness | Middle thickness | Tip thickness | Midrib base thickness | Midrib middle thickness | Midrib tip thickness | Mean thickness | Mean midrib thickness |
| --- | --- | --- | --- | --- | --- | --- | --- | --- | --- | --- |
| Leaf width | **0.489** |  |  |  |  |  |  |  |  |  |
| Base thickness | **0.541** | **0.659** |  |  |  |  |  |  |  |  |
| Middle thickness | **0.206** | *0.154* | **0.570** |  |  |  |  |  |  |  |
| Tip thickness | 0.115 | **0.305** | **0.536** | **0.429** |  |  |  |  |  |  |
| Midrib base thickness | **0.579** | **0.553** | **0.687** | **0.315** | **0.414** |  |  |  |  |  |
| Midrib middle thickness | **0.518** | **0.388** | **0.698** | **0.666** | **0.447** | **0.538** |  |  |  |  |
| Midrib tip thickness | 0.078 | *0.177* | **0.451** | **0.574** | **0.652** | **0.300** | **0.460** |  |  |  |
| Mean thickness | **0.388** | **0.488** | **0.894** | **0.808** | **0.752** | **0.599** | **0.750** | **0.655** |  |  |
| Mean midrib thickness | **0.588** | **0.551** | **0.777** | **0.505** | **0.536** | **0.951** | **0.743** | **0.509** | **0.756** |  |
| L×W | **0.876** | **0.831** | **0.698** | **0.227** | **0.234** | **0.657** | **0.567** | *0.148* | **0.513** | **0.672** |

For this analysis, dimensions of 202 leaves harvested throughout the season were measured. Leaf width: maximum width along the leaf; L×W: length × maximum width. Italic, and bolded values show the significant ( $0.001 < P < 0.05$ ) and very significant correlations ( $P < 0.001$ ), respectively.

**Figure 7.**
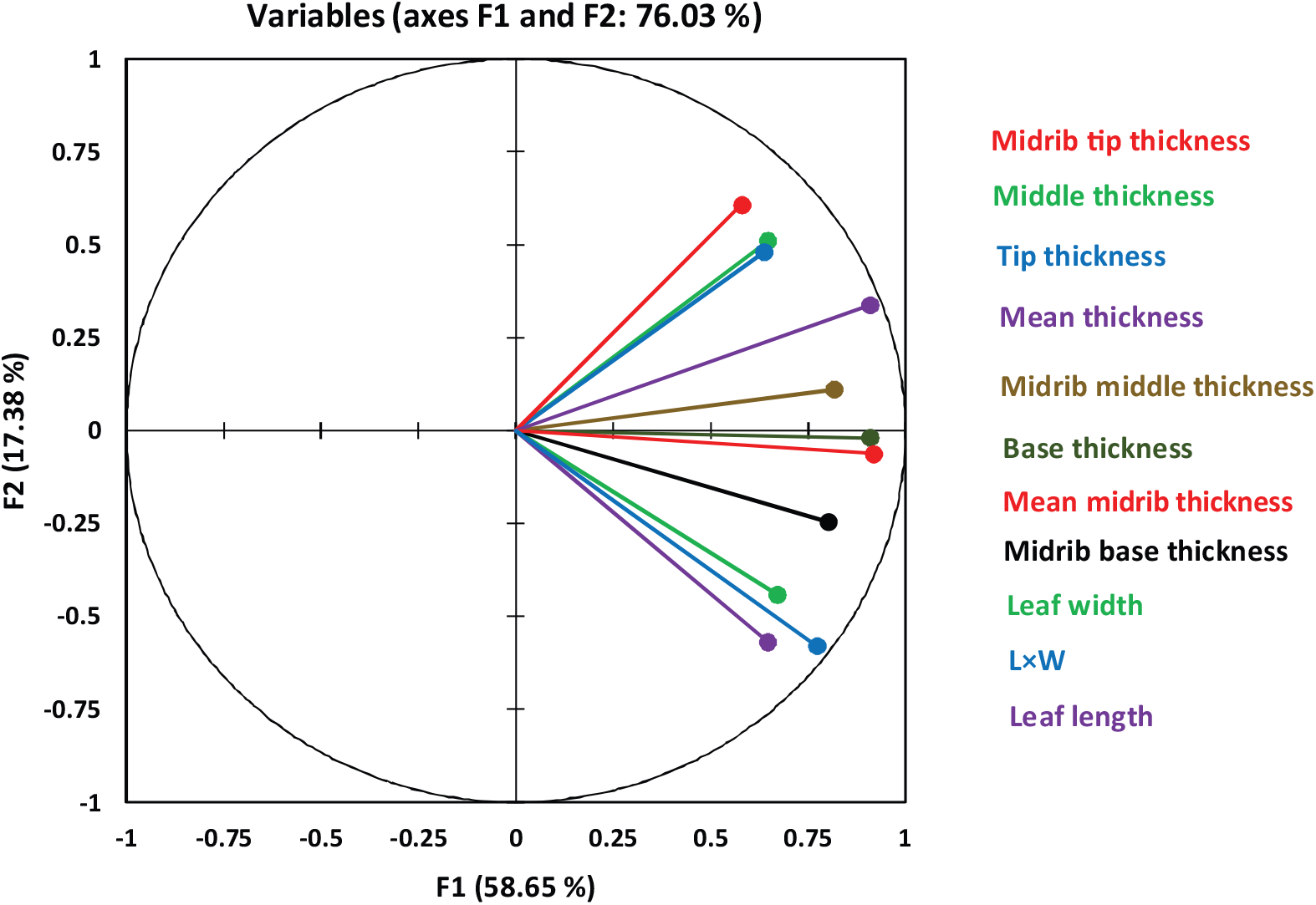
Principal Component Analysis (PCA) of leaf and midrib dimensions. L×W: leaf length × maximum leaf width. Tip, middle and base: thickness at the tip, middle, and base of the leaf.

In the case of using Specific Gravity Bench, the practice of measuring pile leaf area, including the pile volumetry and also determining mean thickness of 10 leaf samples, lasted about 6 minutes (350 seconds, on average; data not shown). Weighing of each leaf pile out of water was done in about 20 seconds. Sampling and reading of one thickness value per leaf (i.e. in the middle of length) also needed about 15 seconds. Besides, volumetry of each pile with 3 replicates, required at most 3 minutes. Remarkably, only 2 or 3 times of repetition were enough to reach a constant value of underwater weight of the immersed set, which highly depended on the formation of bubbles, due to the speed of immersion. The total duration of the practice could be shortened to less than 5 minutes (275 seconds), if 5 leaves were sampled for thickness measurement.

## 1. Discussion

The present study was conducted to test the concept of volumetric leaf area for accelerating leaf area measurements. In the current optical methods, all leaves of a sample should be completely unfolded one by one, and be fed into the imaging or scanning units, separately. Therefore, it is expected that removing this bottleneck may considerably improve the speed of the measurements, provided that the precision of the method is kept at the same level. The idea of volumetric leaf area was calculating the area simply by dividing the volume of leaf sample by its mean thickness. According to the currently available techniques of volumetry and thickness measurement, here the total volume of the pile was measured directly, whereas the mean thickness was estimated by statistical sampling.

Based on the physics principles, as the 3D shape of an object changes, its weight (mass) and volume remain constant. So, the volume of a twisted, folded, or completely flattened leaf is identical. Based on this fact, in the VLA approach, the leaf sample is supposed as a completely flat structure with a known volume and thickness. Hence, regardless of its 2D shape (i.e. the shape of the flat leaf), the volume is simply equal to the 2D area multiplied by thickness. Accordingly, the area can be calculated by dividing the leaf volume by its mean thickness, though, the volume has been measured when the leaf was not really flattened.

Since the values of volumetric leaf area had to be compared with a reference set, i.e. the optically measured area, a brief assessment was conducted on the variations in the scanning resolution. It was observed that the image resolution had not any considerable effect on the precision of the optically measurement of leaf area; hence, even the minimum resolution of 200 dpi (i.e. the images with the largest pixel size) could provide a reliable optical reference for validating the data of volumetric leaf area. Besides, this observation may also provide useful implementations for the optical methods of leaf area measurement, *per se*. Indeed, as there is an inevitable tradeoff between image resolution and speed of scanning, the statistically independence of measurement precision from resolution may provide the opportunity of using the fastest scanners for leaf area measurement. Accordingly, the currently available super-fast scanners with the speed of scanning of an A4 or even A3 size surface in few seconds, although with 200 dpi resolution, may be utilized as a reliable optical device for leaf area measurements.

High correlations between the OLA and VLA values, indicated that the concept of volumetric leaf area could be used as a precise alternative for optically leaf area measurement. Considering that the 3D structure of midrib does not follow the flat shape of the leaf (lamina), the effect of exclusion of midrib volume on the precision of leaf volumetry was also tested. Subtracting the volume of midrib from the leaf volume, almost had not any effect on the precision of prediction (see NRMSEs in Fig. 3); however, it improved the similarity of VLA to OLA by about 5% (see the slopes of the regressed line). Also, the results of Pile Analysis showed a similar trend. Therefore, it can be concluded that regardless of the inclusion or exclusion of the midrib volume in the volumetric calculations, the error of leaf area measurement using the VLA approach was at most 2.61% (see NRMSEs in Figs. 3&4). Such level of error seems to be acceptable comparing with the potential errors occurs in the current optical systems, e.g. due to the folded or overlapped leaves, view angle, lens distortion, etc. Particularly, the advantage of VLA becomes more bolded when the factor of time is included in the comparisons, as a considerable pile of leaves can be assessed simultaneously in a short time, without the need for unfolding or feeding the leaves into the optical units, separately. Consequently, the advantage of utilizing VLA technique enhances as the sample size increases; because the time (and usually the equipment) required for volumetry of a single leaf, and a pile of leaves which weighs hundreds/ or kilos of grams, are almost identical.

The volumetric methods utilized in the present study, i.e. the versions of hydrostatic weighing, are among the most precise, simple, and highly available techniques. The comparative precision of the method in volumetry of small objects is evidenced by Hughes (2005). These methods only require a weighing balance, which is available almost in every laboratory. Moreover, as the type of the required parameters are weight and temperature, identical readings can be recorded by different users; despite some other methods e.g. which are based on monitoring liquid displacement or overflow. Currently, Specific Gravity Benches are widely used in civil engineering, and therefore are readily available in the market of laboratory equipment. One of the advantageous of using SGBs over the original suspension methods, is that there is no need to place the whole system (including water container) on the weighing balance; and thus, the weighing range is limited to the weight of the leaf sample. This option particularly makes it possible to use a more precise balance (or load cell), instead of utilizing the sensors with a wider range of weighing to include the weight of water; as these two properties of balances are often in contrast with each other. Furthermore, in order to facilitate measurement of a higher number of leaf samples, SGBs can become automated and/ or motorized; e.g. for self-recording of weights or water temperature, or having a more controlled and simplified movement of the water container. In general, it is expected that utilizing novel techniques, the volumetric methods will be improved and become even more facilitated and simplified in the future (for instance, see the idea of building a precise acoustic volumeter reported by Sydoruk *et al*., 2020).

Although there may be exclusive techniques or tools for rapid and simultaneous measurement of thickness of multiple objects in the industry, here the only available option was determining the leaf thickness based on the statistical sampling, which is also the accepted cornerstone of almost every evaluation in the biological science. As the results of virtual sampling of 5 leaves with 100 replications indicated (Fig. 5), the actual thickness of population could be predicted in average with less than 2% difference. Also as shown in Table 1, leaf thickness had the least seasonal or diurnal variations among the leaf dimensions (i.e. in comparison with leaf length or width). These observations might be considered as the evidences for the reliability of sampling for leaf thickness estimations. However, compared to the leaf volumetry, in which the volume of the total population can be measured easily, thickness measurement seems to be the relative bottleneck of the VLA approach. Therefore, the VLA may be facilitated (and/or accelerated) yet, by further investigation and utilizing more efficient and innovative techniques of thickness measurement (e.g. see Pfeifer et al., 2018).

Another important observation, was the very high correlation between leaf volume and optical area, compared with the volume-thickness relationship (Fig. 6); the feature which also could be expected theoretically based on the leaf geometry. Indeed, the pattern of variation in leaf volume completely follows that of the leaf area. This characteristic provides two vital opportunities: (i) comparing and evaluating the relative leaf area of various treatments independent of thickness; and (ii) developing a robust simple linear model for leaf area estimation. Indeed, the aim of many leaf area evaluations in plant/ crop science is comparing various genotypes, treatments, or phenological phases, rather than calculating the absolute values of leaf area *per se*. In such cases, assessment of variations in leaf volume may also reflect the changes in the relative leaf area with high degrees of precision and reliability; therefore, there is no need to measure the leaf thickness for converting the sample volume into area. This may be also a computational solution for the bottleneck mentioned before, i.e. the challenges in the thickness measurement. Besides, even where the absolute values of leaf area are required, a robust linear model which estimates leaf area based on the volume may be used; though developing such a general model preferably needs using big data of leaf thickness sampled from various genotypes, phenological phases, and locations. Although this was not among the purposes of the present study, an example of such model is represented only for instance in Fig. 6. Here, the slope of the regression line (around 4), equals to the reverse of mean leaf thickness (i.e. 0.25 mm^-1^); which considering the equation 5, accords to the expectations. Parallel to the concepts of leaf area and LAI, the application of leaf volume may be expanded in the various branches of crop science, e.g. in phenotyping, in the studies of canopy biophysical characteristics, or development of radiative transfer models; as the leaf volume seems to be a more direct contributor to light extinction, compared with leaf area. Thus, the concept of leaf volume may become more widespread and be used interchangeably with leaf area.

In summary, it is expected that utilizing the volumetric leaf area which can be determined faster than the conventional optical area, and also requires simple and available tools, may facilitate the time consuming and laborious practice of destructive leaf area measurement. Consequently, an opportunity may be provided for increasing the number and/ or size of sampling; which in turn, can improve the precision of the field experiments.

## 5. Conclusion

Although nowadays a variety of advanced optical tools are available for destructive leaf area measurement, an important challenge has been remained unsolved, i.e. the need for the time-consuming practice of unfolding the leaves and feeding them into the imaging/ scanning unit, separately. In the present study, the concept of volumetric leaf area was introduced, and its practical application for facilitating leaf area measurement was tested. According to this approach, the leaf area can be calculated simply through dividing the volume of a leaf pile by the mean leaf thickness. It was observed that regardless of the inclusion or exclusion of the midrib volume in the calculations, the VLA values had an approximately 1:1 correlation with the optically measured leaf areas; though neglecting the midrib volume improved the estimations by about 5%. Considering the availability of Specific Gravity Benches, the efficiency of utilizing these tools for volumetry of leaf piles was also evaluated; by which the measurement of each leaf sample (i.e. pile) lasted about 5 to 6 minutes, depended on the number of leaves sampled for determination of thickness.

Furthermore, as it was evidenced that the variations in the leaf area completely follows the pattern of variations of the leaf volume, it is suggested to compare the treatments or samples based on their relative volumes; instead of using the absolute values of leaf area. Therefore, the assessments may become independent of leaf thickness; which can increase the simplicity and the rate of measurements.

Besides the various aspects of VLA, the effect of image resolution on the OLA was also studied, which revealed that there was not any considerable difference between the results of scanning with 200 dpi and 1200 dpi. Consequently, it was suggested that even the superfast scanners with a resolution as low as 200 dpi can be used for optical leaf area measurements.

Considering the results of the present study, it is expected that utilizing the reliable, rapid, and simple technique of volumetric leaf area, in which the required measurement time may be independent of the sample size, facilitate the laborious practice of destructive leaf area measurement; and consequently, improve the precision of field experiments.

